# Phenotype Dependent Segregation of a Novel *EPS8* Variant for Hearing Loss and an *HPDL* Variant for Neurodevelopmental Disorders in a Complex Consanguineous Family

**DOI:** 10.64898/2026.07.09.737440

**Authors:** Bassam Jamalalail, Ahmed Khalifa, Bipin Balan, Shuhd Bineshaq, Dia Advani, Hanan Elsokary, Kondaramage Dasuki, Suhana Shiyas, Nelson C. Soares, Shehzad Hanif, Samuel Mathew Tharakan, Zeinab Mohamdi, Mohammad Rajeh Aburaidah, Sherin Kuttiankandy, Alawi Alsheikh-Ali, Nasna Nassir, Mohamad El Bitar, Mohammed Uddin

## Abstract

Consanguinity increases the risk of autosomal recessive disorders and may result in the co-segregation of multiple pathogenic variants within the same family. Although most affected families are explained by a single genetic diagnosis, multilocus pathogenic variation can produce complex and overlapping clinical phenotypes. We investigated a consanguineous Pakistani family with three affected siblings, including dizygotic twins presenting with neurodevelopmental disorder and hearing loss, using detailed clinical evaluation, long-read whole-genome sequencing, bulk transcriptomics, protein profiling, and segregation analysis to determine the underlying molecular diagnoses. One sibling presented with isolated non-syndromic hearing loss, whereas the dizygotic twins exhibited severe neurodevelopmental impairment characterized by global developmental delay, spastic quadriplegic cerebral palsy, microcephaly, and white matter abnormalities. Long-read whole-genome sequencing identified a homozygous start-loss variant in *HPDL* (c.3G>C) in both twins, consistent with *HPDL*-related neurodevelopmental disorder with progressive spasticity and brain white matter abnormalities (NEDSWMA). In addition, a novel homozygous nonsense variant in *EPS8* (c.1294C>T) was identified in one twin and the sibling with isolated hearing loss, explaining the auditory phenotype. Long-read transcriptomic analysis demonstrated absence of detectable *EPS8* transcripts in both individuals homozygous for the nonsense variant, providing transcript-level evidence consistent with a loss-of-function mechanism. Genome-wide comprehensive proteomic profiling (SomaScan) identified distinct protein abundance profiles across family members, with the most pronounced alterations observed in the twins affected by *HPDL*-related neurodevelopmental disease, particularly the individual harboring pathogenic variants in both *EPS8* and *HPDL*. This study expands the mutational spectrum of *EPS8* and highlights the independent segregation of two autosomal recessive disorders within the complex consanguineous family, resulting in distinct and blended phenotypes.

## Introduction

Consanguinity is defined as the union of individuals who are related as second cousins or closer (1). Consanguineous marriages have been a longstanding cultural tradition in many populations across North Africa, the Middle East, and West Asia, where unions between biological relatives account for approximately 20% to more than 70% of all marriages (2). Pakistan is among the countries with the highest prevalence of consanguineous marriages worldwide, with cousin unions accounting for approximately 65% of all marriages (3,4). Pakistan, a culturally diverse nation characterized by complex ethnic and caste structures, consistently reports one of the highest rates of consanguineous marriage worldwide (4–6). Consanguinity has important health and reproductive implications. The increased sharing of genetic material (i.e. runs of homozygosity) between related partners elevates the risk of homozygosity for deleterious recessive variants, thereby increasing the likelihood of inherited genetic disorders (4,7–9). Numerous studies have linked consanguineous marriages to adverse reproductive and pregnancy outcomes, including higher rates of miscarriage, stillbirth, pregnancy termination, low birth weight, congenital anomalies, and infant mortality (7,10–15). Therefore, while consanguineous marriages remain socially and culturally valued in Pakistan, they also contribute to a greater burden of genetic and reproductive health risks (4). Although inherited disorders in consanguineous families are mostly reported to have a single autosomal recessive pathogenic variant, disease causation may not always be attributable to a single locus for a family. In highly consanguineous or endogamous populations, multiple rare recessive variants can segregate within the same extended pedigree, resulting in distinct Mendelian disorders among affected relatives or blended phenotypes in individuals harboring pathogenic variants at more than one gene. Consequently, multilocus inheritance should be considered when the clinical presentation is unusually complex or cannot be fully explained by a single molecular diagnosis (16–19)

Hearing loss is the most common sensory impairment in newborns, affecting approximately 1.33 per 1000 live births (20). Hereditary hearing loss is broadly categorized into syndromic and non-syndromic forms. Syndromic hearing loss is accompanied by additional clinical manifestations affecting other organ systems, whereas non-syndromic hearing loss (NSHL) is characterized by isolated auditory impairment. NSHL accounts for nearly 70% of inherited hearing loss, while the remaining 30% is syndromic. Among individuals with NSHL, autosomal recessive inheritance is the predominant pattern, representing approximately 75–80% of cases, whereas autosomal dominant, X-linked, and mitochondrial forms occur less frequently (21–25). The molecular basis of NSHL is remarkably heterogeneous, involving pathogenic variants in more than a hundred genes. Although variants in genes such as *GJB2, SLC26A4, OTOF*, and *TMC1* account for a substantial proportion of genetically diagnosed cases, many less frequently implicated genes continue to be identified, highlighting the extensive genetic diversity underlying this condition (26,27).

Neurodevelopmental disorders (NDDs) comprise a spectrum of rare conditions arising from disruptions in brain development that impair neurological function. These disorders are characterized by varying degrees of deficits and comorbidities in cognition, motor skills, communication, and adaptive function, reflecting their broad clinical and genetic heterogeneity (28). The etiology of NDDs is multifactorial, with both genetic and environmental factors contributing to disease development (29).

Here, we report a consanguineous family in which multiple autosomal recessive disorders segregate independently. The family includes one child with isolated non-syndromic hearing loss and dizygotic twin siblings affected by a severe NDD caused by *HPDL* variants, one of whom also presents with hearing loss. This family highlights the complexity of genetic diagnosis in consanguineous pedigrees and underscores the importance of comprehensive genomic analysis of full family for identifying multilocus pathogenic variation.

### Phenotypic Presentation

We investigated a consanguineous family of Pakistani origin comprising healthy parents and four offspring, including one dizygotic twin (male and female) (Figure 1). The family was referred for genetic evaluation because of congenital hearing loss in two siblings and a severe NDD in dizygotic twin siblings. The parents are first cousins with reported history of hearing loss and neurodevelopmental disorders. Three children were affected by inherited disorders with variable clinical presentation. One sibling presented with isolated bilateral sensorineural hearing loss, where the dizygotic twin siblings exhibited severe neurodevelopmental impairment. Moreover, one of the twins also has hearing loss, suggesting the coexistence of two distinct phenotypes within the same family.

**Figure 1.**
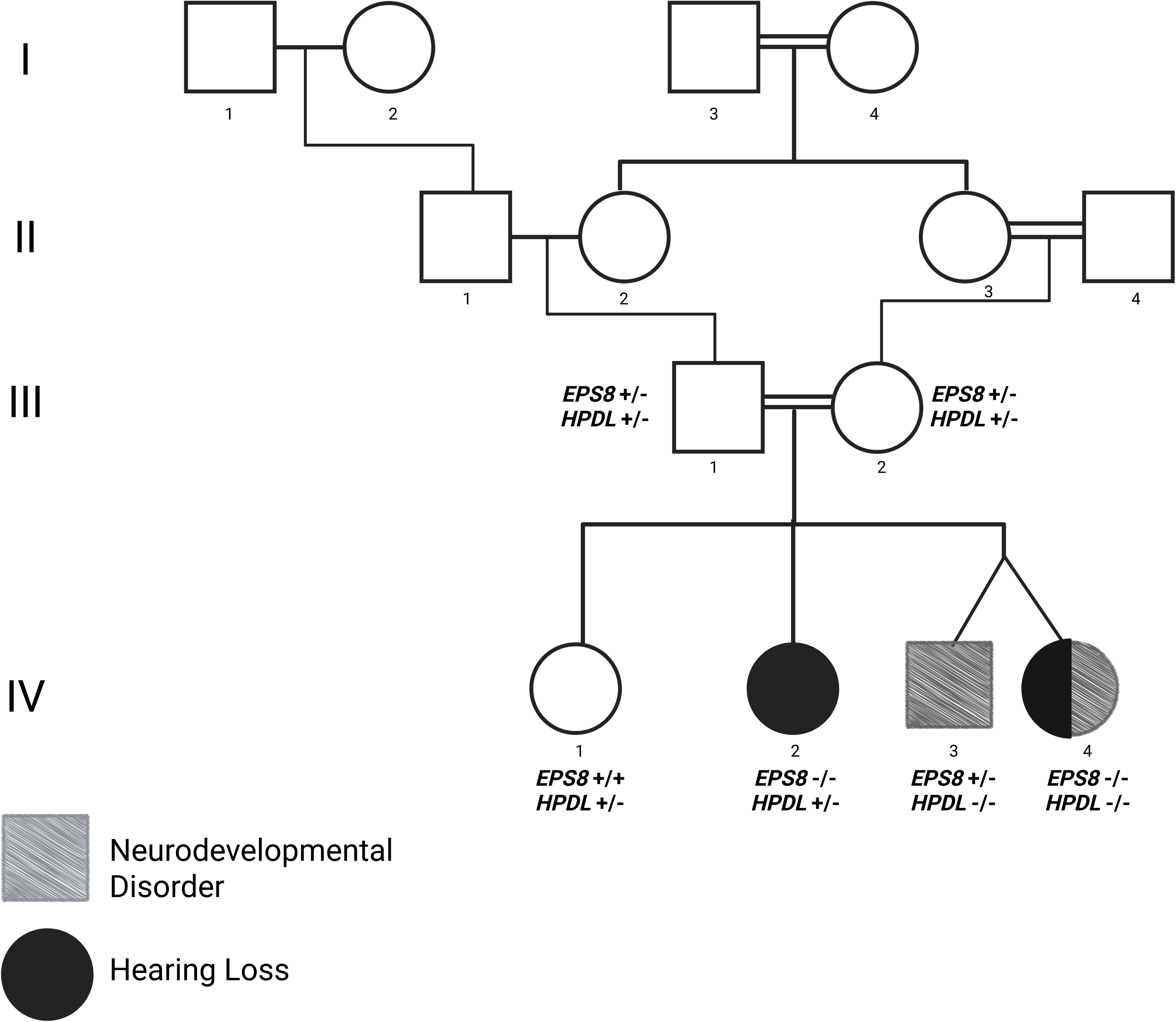
Pedigree and segregation of hearing loss and neurodevelopmental disorder in the studied consanguineous Pakistani family. Pedigree of the consanguineous Pakistani family demonstrating the segregation of two independently inherited autosomal recessive disorders. Squares represent males and circles represent females. Double horizontal lines indicate consanguineous unions. Filled black symbols denote individuals affected with non-syndromic hearing loss, while hatched gray symbols denote individuals affected with neurodevelopmental disorder with progressive spasticity and brain white matter abnormalities (NEDSWMA). The half-filled symbol represents an individual affected with both hearing loss and NEDSWMA. Genotypes for the identified *EPS8* (NM_004447.5:c.1294C>T; p.Arg432Ter) and *HPDL* (c.3G>C; p.(Met1?)) variants are shown beneath each tested individual. For each gene, +/+ indicates the homozygous reference genotype, +/− indicates the heterozygous carrier genotype, and −/− indicates the homozygous pathogenic genotype. The pedigree demonstrates independent segregation of the novel EPS8 nonsense variant with hearing loss and the HPDL start-loss variant with NEDSWMA, with the female dizygotic twin harboring homozygous pathogenic variants in both genes, resulting in a blended phenotype.

## Methods

### Subjects

The family was referred to the Center for Applied and Translational Genomics (CATG) at Mohammed Bin Rashid University of Medicine and Health Sciences (MBRU) for inclusion in our Rare Genetic Diseases project, which provides long-read genome sequencing (LR-WGS) to patients with suspected rare genetic conditions. They were enrolled due to the proband’s hearing loss as well as the presence of a NDD in two siblings. The research activity conducted in this study complied with applicable ethical standards. The study received approval from the Institutional Review Boards of the Dubai Scientific Research Ethics Committee (DSREC-06/2025_11) and Mohammed Bin Rashid University of Medicine and Health Sciences (MBRU-IRB-2023-335).

### Sample Collection and Phenotyping

Peripheral blood samples were collected from six members of the affected family, including the proband, both parents, and three siblings, after obtaining written informed consent. Sample collection was carried out at Al Jalila Children’s Hospital under sterile conditions using standard procedures. All samples were labeled with anonymized identifiers to ensure confidentiality and were stored at –80°C until further analysis

### DNA Isolation and Sequencing

#### Pacific Biosciences High-Fidelity Sequencing

High molecular weigh genomic DNA was extracted from 200uL of flash-frozen blood samples using the GenFind V3 (Cat. No. C34881) and Nanobind DNA (Cat. Np. 102-762-700) extraction kits, following the manufacturer’s protocols. DNA concentration was quantified using a Qubit 3 fluorometer (Thermo Fisher, USA) in conjunction with dsDNA HS Assay Kit (Cat. No. Q33231), and samples were diluted to 30ng/uL in 30uL volumes for subsequent shearing. Shearing was performed using the Diagenode Megarupter 3 (Belgium) with hydropore-syringes (Cat. No. E07010003), targeting fragment sizes of 15–18 kb, as per PacBio’s guidelines. Fragment lengths were assessed using the Agilent 4200 TapeStation and genomic DNA screen tape (Cat. No. 5067-5365). SMRTbell libraries were prepared using the SMRTbell Prep Kit 3.0 (Cat. No. 102-182-700). Sequencing was performed on the PacBio Revio platform (Pacific Biosciences, Menlo Park, CA) using Revio SMRT Cells (Cat. No. 102-202-200) and the corresponding polymerase kit (Cat. No. 102-817-600). The run protocol incorporated adaptive loading, with a 2-hour pre-extension and 30-hour movie acquisition.

#### Bulk Transcriptome Sequencing

Total RNA was extracted from samples obtained from all six family members using the RNeasy Mini Kit (Qiagen) according to the manufacturer’s instructions. Messenger RNA (mRNA) was isolated using the Dynabeads™ mRNA DIRECT™ Purification Kit (Invitrogen). Purified mRNA was reverse transcribed to generate double-stranded cDNA using the Maxima Minus Double-Stranded cDNA Synthesis Kit (Thermo Scientific). The resulting cDNA was purified using AMPure XP beads (Beckman Coulter) and used for library preparation with the Oxford Nanopore Technologies Ligation Sequencing Kit (LSK-109) according to the manufacturer’s protocol. Individual samples were barcoded prior to library preparation and sequenced on the Oxford Nanopore PromethION platform using R9.4.1 flow cells (FLO-PRO002).

#### Protein Profiling

Relative protein abundance in plasma samples obtained from the study participants was quantified using the SomaScan® platform (SomaLogic Inc., Boulder, CO, USA) (30), an aptamer-based proteomic assay that employs a library of 11,000 Slow Off-rate Modified Aptamers (SOMAmers) to selectively bind target proteins. Protein abundance was measured by DNA microarray and reported as relative fluorescence units (RFUs). Quality control and normalization were performed according to SomaLogic’s standard analytical workflow, which incorporates multiple internal calibrators and quality control samples to ensure assay performance. Principal component analysis (PCA) of log₂-transformed protein abundance data was performed at both the sample and aptamer levels to identify potential outliers. No additional samples or aptamers were excluded following PCA. Assay reproducibility was assessed using the coefficient of variation (CV) of calibrator and quality control samples, with all plates meeting the recommended quality thresholds. Aptamers targeting non-human proteins, non-protein molecules, or lacking corresponding gene annotations were excluded prior to statistical analyses.

#### Bioinformatics & Variant Analysis

The quality of PacBio HiFi long reads was evaluated using Cramino (v1.0.0) (31). Read alignment to the Homo sapiens reference genome (GRCh38) was performed with pbmm2 (v1.17) (32), and the resulting BAM files were sorted using Samtools (v1.21) (33). DeepVariant (v1.9.0) (34) was employed for single nucleotide variant (SNV) and small indel calling. Phasing of aligned BAM and VCF files was carried out with WhatsHap (v2.7) (35). Structural variants were identified using PBSV (v2.11) (36).

The variant annotation of the passed PacBio variants was done by ANNOVAR (version 2020-06-08) (37) based on the reference genome of GRCh38. The following databases were used for ANNOVAR annotations: refGene, cytoBand, dbnsfp42a, avsnp150, clinvar_20240611, gene4denovo201907, and gnomad41_genome. The following filters were further applied to identify the variants: (i) Mutations with depth <8 were discarded. (ii) Mutations listed in gnomAD allele frequency >0.001 were excluded (iii) Mutations listed in the Clinvar having values ’Benign’ and ’protective’ were removed and (iv) considered only the variants which are in ’exonic’,’splicing’ or ’exonic;splicing’. From the PBSV VCF file, we considered only structural variants (SVs) ≥50 bp for the downstream analysis. Additionally, we used multiple variant annotation and classification tools according to American College of Medical Genetics and Genomics (ACMG) guidelines (38). This includes ANNOVAR and Horizon 2.3.0, which predicts ACMG guided pathogenicity and allows integration of a custom clinical genetic variant database with other large-scale publicly available databases (e.g., gnomAD, ClinVar, pLI, AM, CADD). Variant classifications were cross checked with both tools for consensus pathogenicity (38). We used Horizon to infer SV pathogenicity, as it supports pathogenicity prediction for long-read-based structural variants (39).

Long-read transcriptome sequencing was performed on RNA obtained from all six members of the nuclear family. Raw reads were quality-filtered using NanoFilt (minimum quality score Q10) and aligned to the human transcriptome reference using minimap2 in splice-aware long-read mode. Gene-level transcript abundance was quantified using NanoCount, and merged gene-count tables were generated for each individual. Gene annotation was performed using the project-specific reference annotation to ensure consistency with the transcriptomic reference. Gene counts were normalized using edgeR, converted to log counts per million (logCPM), and transformed into gene-wise Z-scores across all six family members. Relative transcript abundance was visualized using a predefined *EPS8*/*HPDL* mutation-relevant pathway panel. In addition to pathway-level analysis, transcript detection of *EPS8* was assessed across all family members

An exploratory family-based proteomic analysis was performed using SomaScan v5.0 normalized RFU data. The ADAT file was parsed in Python; protein features were mapped from SomaScan SeqIDs to Entrez gene symbols, and only biological study samples were retained after excluding buffer, QC, and calibrator rows. We defined genotype and phenotype-informed contrasts to compare EPS8 homozygous samples, HPDL homozygous samples, the dual-affected individual, and individual affected siblings against the unaffected sibling. For each protein, I calculated log2 fold change using normalized RFU values with a pseudocount of 1. Mann–Whitney U tests were applied only to contrasts with at least two samples per group; 1-vs-1 comparisons were treated as exploratory fold-change-only contrasts. The visualization focused on a curated EPS8 pathway protein set, including EGFR/RTK, Ras-Rac, kinase, membrane protrusion/cytoskeletal, and HPDL-related proteins. The bar plots show per-contrast log2 fold changes, while the heatmap summarizes directional consistency across all contrasts.

#### Sanger Sequencing

The novel EPS8 nonsense variant, NM_004447.5:c.1294C>T (p.Arg432Ter), identified by long-read whole-genome sequencing was validated by bidirectional Sanger sequencing in all six members of the nuclear family. A single primer pair flanking the target variant (forward: 5′-CCTGTTTGCGCTGATGTTCT-3′; reverse: 5′-AACTTGCTGGGCTGTTTCTC-3′) was designed to amplify a 638-bp PCR product encompassing the variant locus. PCR amplification was performed using an annealing temperature of 60°C, followed by bidirectional sequencing using the BigDye Terminator v3.1 Cycle Sequencing Kit (Applied Biosystems) on an ABI PRISM 3730XL Genetic Analyzer. Sequence chromatograms were analyzed using Variant Reporter Software v2.1 (Applied Biosystems), DNASTAR Lasergene SeqMan 7.0, and the Macrogen SNP Analysis program (v2.0). Sanger sequencing was performed to validate the lrWGS finding and confirm segregation of the EPS8 variant within the family

## Results

### Clinical Findings

We investigated a consanguineous Pakistani family with two distinct phenotypes segregating with the same pedigree (Figure 1). The proband (IV-2), a 13-year-old female, presented with isolated congenital hearing loss. Audiological assessment confirmed bilateral profound sensorineural hearing loss. No additional syndromic features were identified.

The dizygotic twins (IV-3 and IV-4) were born prematurely at 35 weeks’ gestation by emergency cesarean section following premature rupture of membranes and had low birth weights. Both required neonatal intensive care admission for two weeks. Developmental delay became apparent during infancy, and both subsequently developed a severe neurodevelopmental phenotype characterized by spastic quadriplegic cerebral palsy (Gross Motor Function Classification System [GMFCS] Level V), profound global developmental delay, intellectual disability, microcephaly, failure to thrive, and minimal verbal communication. Neither twin achieved independent sitting, standing, or walking, and both required maximal assistance for all activities of daily living.

Neurological examination demonstrated truncal hypotonia with marked spasticity and dystonic posturing affecting all four limbs, tight hip adductors and hamstrings, bilateral ankle contractures, and neuromuscular scoliosis. Orofacial dysfunction with dysphagia was present in the female twin, resulting in feeding difficulties and recurrent chest infections. The male twin additionally exhibited hand stereotypies and oromandibular dystonia.

Brain magnetic resonance imaging performed in the male twin demonstrated generalized cerebral atrophy with reduced white matter volume, hypoplasia of the corpus callosum, mild prominence of the lateral ventricles, and bilateral periventricular white matter hyperintensities on T2-weighted and FLAIR sequenced, consistent with white matter abnormalities. A neonatal brain MRI performed in the female twin was reported to show abnormal white matter changes. Although the twins shared a similar neurodevelopmental phenotype, only the female twin (IV-4) had congenital bilateral profound sensorineural hearing loss confirmed by audiological assessment, whereas hearing evaluation in the male twin (IV-3) was normal. The clinical characteristics and molecular diagnoses of the affected siblings are summarized in (Table 1).

**Table 1.**
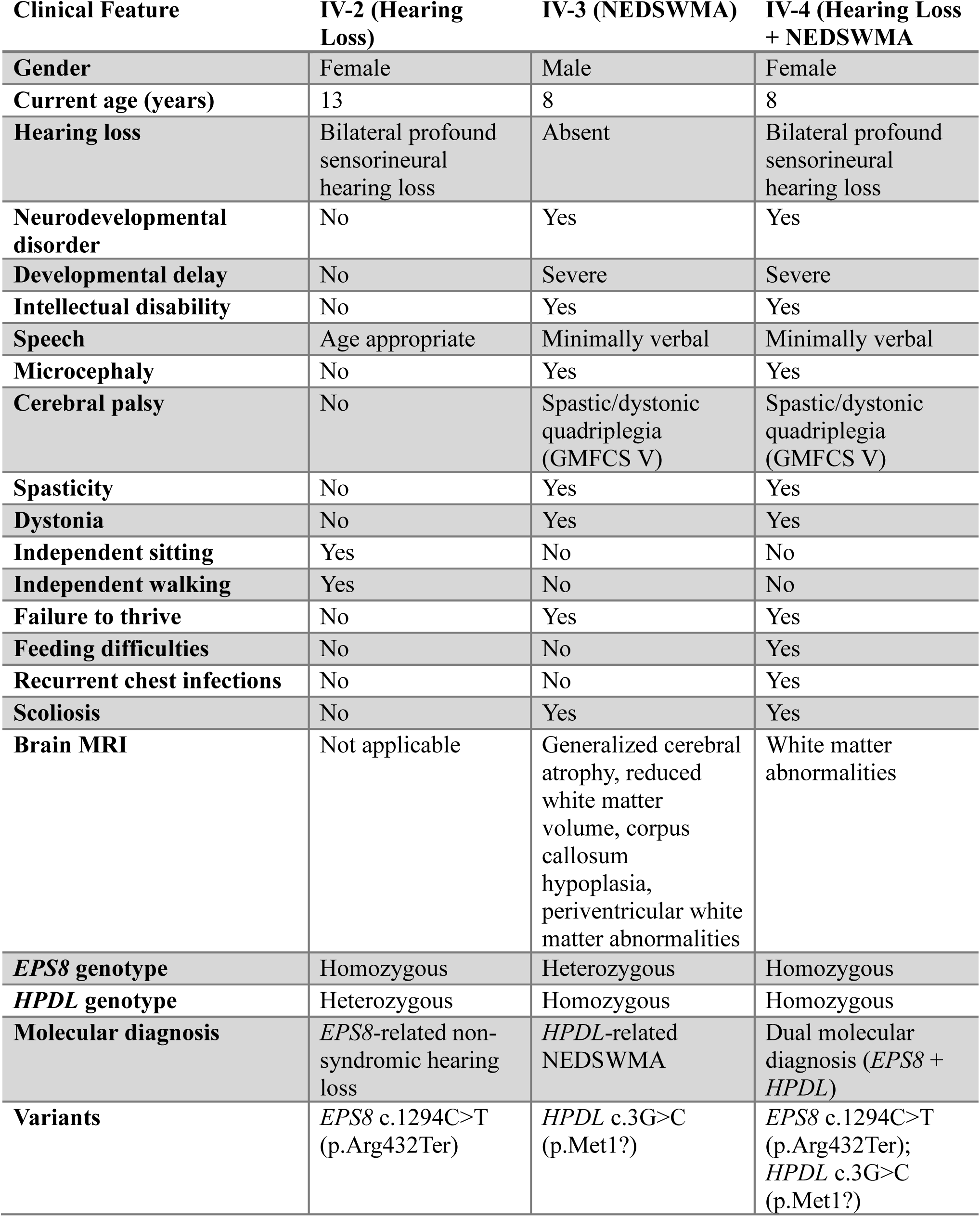
Clinical characteristics of the affected siblings.

### Genetic Findings

Long-read whole-genome sequencing (lrWGS) was performed on all six members of the nuclear family, including both parents and their four children. Analysis identified two independently segregating autosomal recessive disorders within the family (Figure 1).

A homozygous nonsense variant in *EPS8*, 1294C>T; p.Arg432Ter, was identified in the sibling with isolated hearing loss (IV-2). The same homozygous variant was also detected in the female dizygotic twin (IV-4), who presented with hearing loss in addition to neurodevelopmental impairment. Both parents (III-1 and III-2) and the male dizygotic twin (IV-3) were heterozygous carriers, whereas the unaffected sibling (IV-1) was homozygous for the reference allele. The variant and its segregation pattern were subsequently confirmed by Sanger sequencing in all six family members, with complete concordance between the Sanger sequencing and lrWGS results (Supplementary Figure S1). The segregation pattern was consistent with autosomal recessive inheritance and explained the hearing loss phenotype observed in IV-2 and IV-4.

In addition, both dizygotic twins (IV-3 and IV-4) were found to harbor a homozygous start-loss variant in the *HPDL* gene, 3G>C; p.Met1?. Both parents were heterozygous carriers of the variant. The unaffected sibling (IV-1) and the hearing loss sibling (IV-2) were also heterozygous carriers, whereas neither carried biallelic *HPDL* variants. The homozygous *HPDL* variant segregated exclusively with the neurodevelopmental phenotype, establishing the molecular diagnosis of neurodevelopmental disorder with progressive spasticity and brain white matter abnormalities (NEDSWMA) in the twins.

Overall, lrWGS demonstrated the independent segregation of pathogenic variants in EPS8 and *HPDL* within the same consanguineous family. The female dizygotic twin (IV-4) was homozygous for pathogenic variants in both genes, accounting for her blended phenotype of congenital hearing loss and HPDL-related neurodevelopmental disorder. In contrast, her twin brother (IV-3) was homozygous only for the *HPDL* variant and exhibited NEDSWMA without hearing loss, while the sibling with isolated hearing loss (IV-2) was homozygous only for the *EPS8* variant.

### Variant Interpretation

The identified *EPS8* variant, c.1294C>T (p.Arg432Ter), is a novel nonsense variant predicted to introduce a premature termination codon, resulting in loss of normal protein function. To our knowledge, this variant has not been previously reported in the published literature, ClinVar, or disease-specific variant databases. It is also extremely rare in population databases, with only four heterozygous alleles and no homozygous individuals reported in gnomAD. Given the established association of biallelic loss-of-function variants in *EPS8* with autosomal recessive non-syndromic hearing loss, together with its predicted functional consequence and segregation with the hearing loss phenotype in this family, the variant was classified as likely pathogenic according to the ACMG/AMP guidelines based on the PVS1, PM2_Supporting, PP1, and PP4 criteria.

The *HPDL* variant (NM_032756.4:c.3G>C; p.Met1?) affects the canonical translation initiation codon and is predicted to abolish normal translation initiation. Based on the ACMG/AMP guidelines, this variant was also classified as likely pathogenic, supported by the PVS1_Moderate, PM2_Supporting, PP1, and PP4 criteria. The segregation of the identified variants corresponded closely with the observed phenotypes. Biallelic *EPS8* variants were identified only in individuals with hearing loss, whereas biallelic *HPDL* variants were present exclusively in the twins affected by NEDSWMA. The female twin, who was homozygous for likely pathogenic variants in both genes, exhibited the combined phenotype of hearing loss and NDD.

### Transcriptomics Analysis

#### Relative expression profiling

Long-read transcriptome sequencing was performed on all six family members to evaluate transcript abundance of genes within the predefined *EPS8/HPDL* mutation-relevant pathway panel. Gene-wise Z-score analysis demonstrated distinct relative expression patterns among family members across genes involved in EGFR/RTK signaling, Ras-to-Rac actin signaling, kinase regulation of *EPS8*, protrusions and microvilli formation, Wnt signaling/endocytosis, and stereocilia-associated pathways (Figure 2A). As the expression values were normalized across the family, the heatmap represents relative within-family transcript abundance rather than formal differential expression.

**Figure 2.**
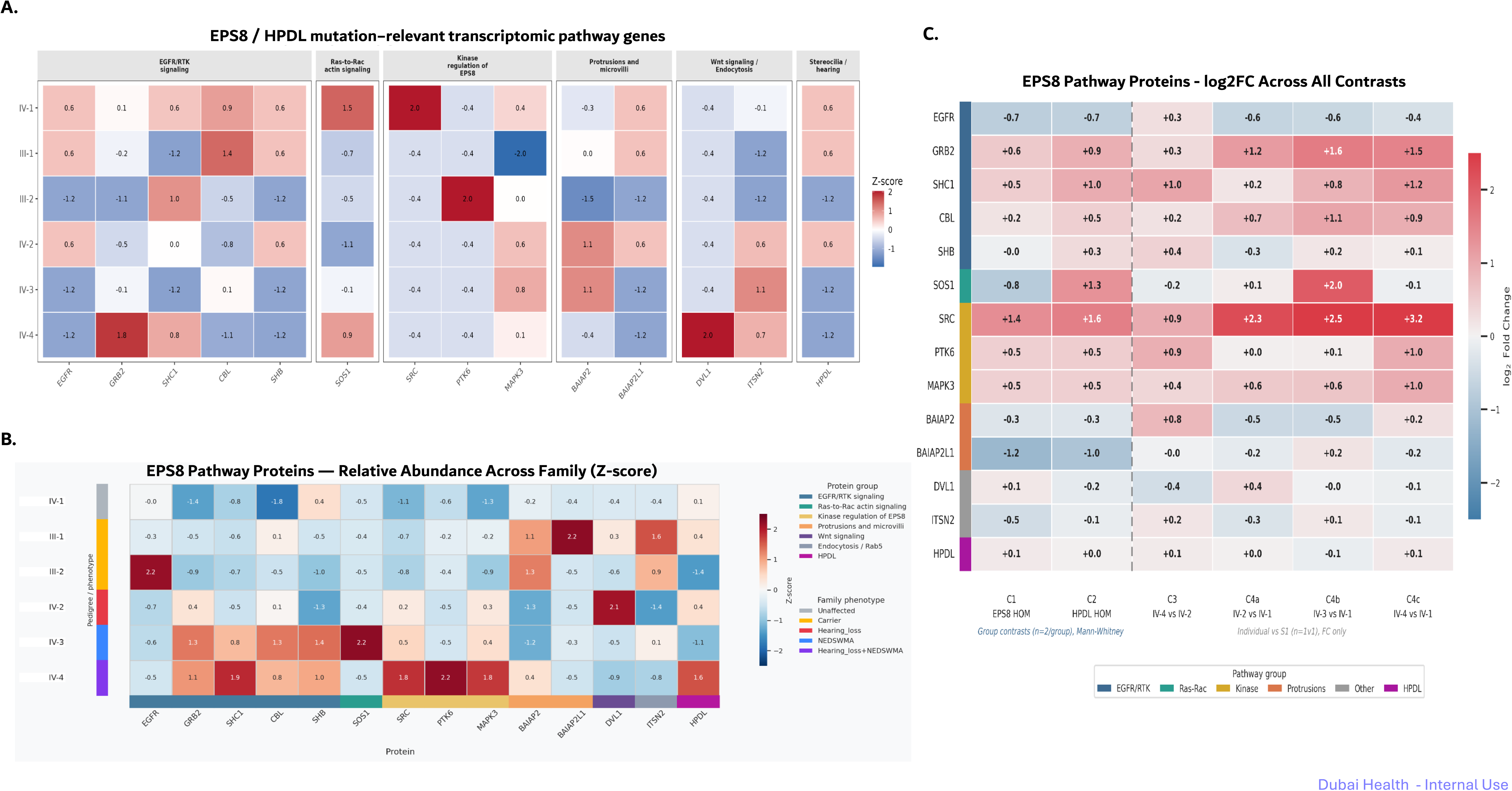
Multi-omics characterization of the EPS8/HPDL mutation-relevant pathway in the studied family. **(A)** Heatmap showing the relative transcript abundance of genes within the predefined EPS8/HPDL mutation-relevant pathway across all six family members. Gene-level counts obtained from long-read transcriptome sequencing were normalized using edgeR, converted to log counts per million (logCPM), and transformed into gene-wise Z-scores across the family. Expression values therefore represent relative within-family transcript abundance rather than formal differential expression. Genes are grouped according to their functional roles, including EGFR/RTK signaling, Ras-to-Rac actin signaling, kinase regulation of EPS8, protrusions and microvilli, Wnt signaling/endocytosis, and stereocilia/hearing. Red indicates relatively higher transcript abundance, whereas blue indicates relatively lower transcript abundance for each gene. **(B)** Heatmap showing the relative abundance of proteins within the predefined EPS8/HPDL mutation-relevant pathway across all six family members. Plasma protein abundance was measured using the SomaScan platform and normalized as gene-wise Z-scores to visualize relative protein abundance within the family. Proteins are grouped according to their functional categories, and the annotation bar indicates the clinical phenotype of each family member. Red represents relatively higher protein abundance, whereas blue represents relatively lower protein abundance. **(C)** Heatmap illustrating pairwise comparisons of protein abundance within the predefined EPS8/HPDL mutation-relevant pathway. Protein abundance is presented as log 2 fold change (log 2FC) across predefined group and individual comparisons. Group comparisons (C1 and C2) were performed using the Mann–Whitney U test (n = 2 per group), whereas individual comparisons (C3–C4c) represent fold-change values relative to the unaffected sibling (IV-1). Positive values (red) indicate relatively increased protein abundance, whereas negative values (blue) indicate relatively decreased protein abundance.

#### Targeted transcript detection

Targeted transcript detection analysis demonstrated detectable *EPS8* transcripts in both heterozygous carrier parents (III-1 and III-2), the unaffected sibling (IV-1), and the sibling heterozygous for the *EPS8* variant (IV-3). In contrast, *EPS8* transcripts were not detected in the two siblings homozygous for the novel nonsense variant (IV-2 and IV-4). *HPDL* transcripts were detected in all six family members irrespective of genotype. The concordant findings obtained using transcript-level and gene-level counting approaches support loss of detectable *EPS8* transcript in individuals homozygous for the c.1294C>T (p.Arg432Ter) variant.

#### Proteomics Analysis

Proteomic profiling was performed on plasma samples obtained from all six family members to investigate alterations in the EPS8 signaling pathway. Relative protein abundance was assessed across proteins functionally related to EPS8 and compared between affected and unaffected family members. The individual-level analysis demonstrated distinct protein abundance profiles among family members (Figure @2). The individual harboring the dual molecular diagnosis (IV-4) exhibited the greatest perturbation of the EPS8 pathway, with increased abundance of several signaling proteins, including SHC1, SRC, PTK6, MAPK3, and GRB2, compared with unaffected family members. The sibling with isolated HPDL-related NDD (IV-3) showed a more modest pattern of pathway dysregulation, whereas the sibling with isolated hearing loss (IV-2) demonstrated relatively limited changes in the EPS8 signaling network. The heterozygous carrier parents also displayed distinct protein abundances despite the absence of clinical manifestations (Figure 3).

**Figure 3.**
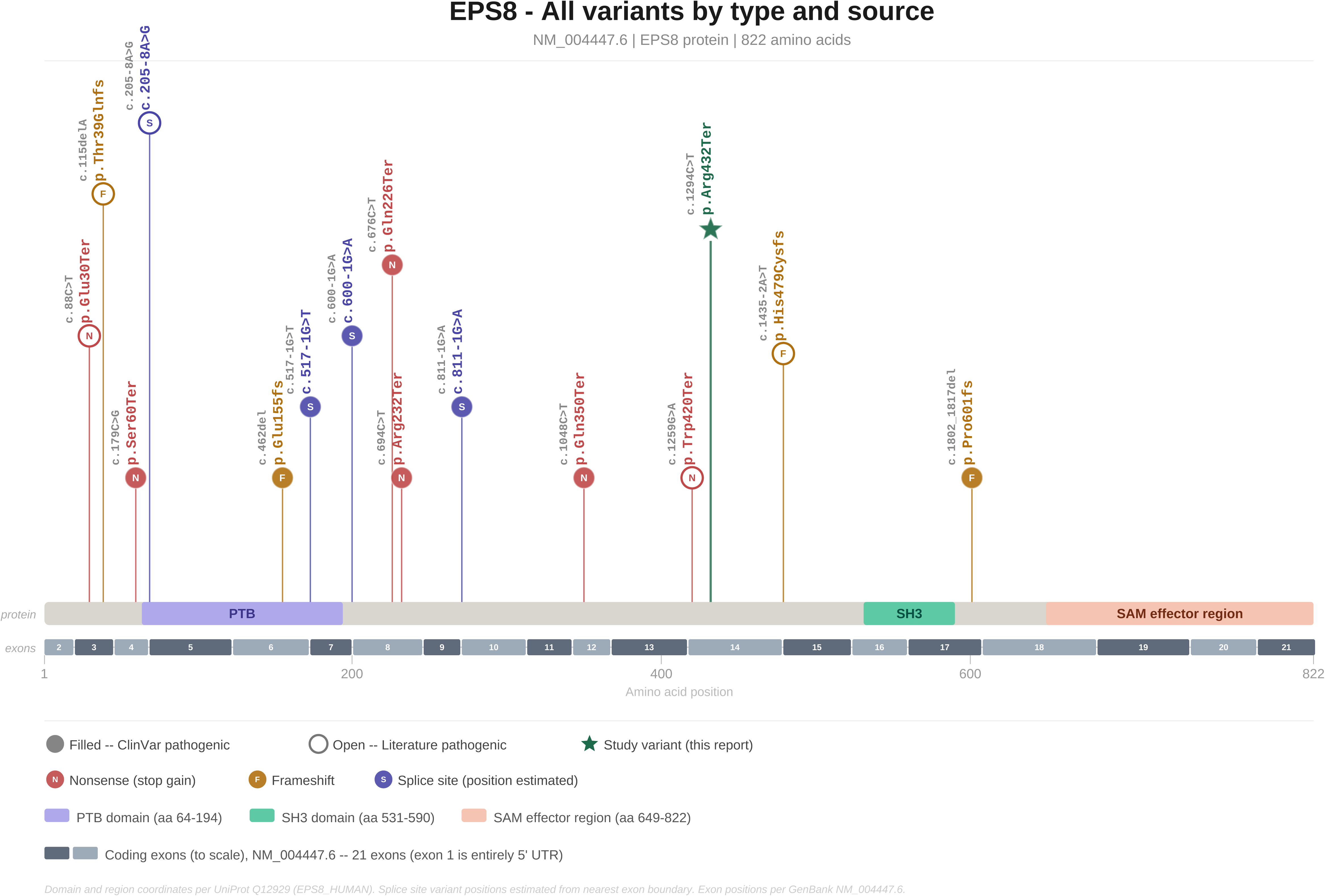
Distribution of reported pathogenic EPS8 variants highlighting the novel c.1294C>T (p.Arg432Ter) variant identified in this study. Schematic representation of the EPS8 protein showing the location of previously reported pathogenic variants together with the novel nonsense variant identified in this study. Variants reported in ClinVar and the literature are displayed according to their predicted consequence (nonsense, frameshift, and splice-site variants). The novel EPS8 c.1294C>T (p.Arg432Ter) variant identified in the present family is indicated by the green star. Protein domains, including the PTB domain, SH3 domain, and SAM effector region, are shown beneath the protein schematic. The distribution of pathogenic variants across the coding sequence supports loss of function as the principal disease mechanism underlying autosomal recessive EPS8-associated hearing loss

To further explore pathway-level differences, relative protein abundance was examined across multiple predefined pairwise comparisons (Figure 4). Comparison of individuals homozygous for the *EPS8* variant with those homozygous for the *HPDL* variant showed higher relative abundance of several proteins within the predefined EPS8/HPDL mutation-relevant panel, including SRC, SHC1, GRB2, CBL, PTK6, and MAPK3, with SRC exhibiting the largest fold changes. Similar expression patterns were observed when the individual with the dual molecular diagnosis was compared with the unaffected sibling. These findings suggest consistent differences in the abundance of selected proteins within the predefined pathway panel across affected individuals. However, given that these analyses were performed in plasma samples from a single family with a limited sample size, including individual (1-versus-1) comparisons, the proteomic findings should be considered exploratory and hypothesis-generating rather than evidence of pathway activation or direct functional effects. Proteins outside the core signaling panel generally exhibited smaller relative differences across comparisons.

Collectively, these findings demonstrate differential regulation of proteins within the *EPS8*-associated signaling network across family members and support a functional effect of the novel *EPS8* nonsense variant. The most pronounced proteomic alterations were observed in the individual harboring pathogenic variants in both *EPS8* and *HPDL*, suggesting that the combined molecular diagnoses may contribute to a more extensive perturbation of downstream signaling pathways.

## Discussion

Consanguineous unions increase the probability of homozygosity for rare pathogenic variants inherited from a common ancestor and are therefore associated with an increased burden of autosomal recessive disorders. Although a single molecular diagnosis accounts for most affected families, highly consanguineous pedigrees may segregate multiple recessive disorders simultaneously, resulting in blended or apparently atypical phenotypes (40–42). In this study, we describe a consanguineous Pakistani family with two independently segregating autosomal recessive disorders caused by likely pathogenic variants in the *EPS8* and *HPDL*. LrWGS identified a novel homozygous nonsense variant in *EPS8* (c.1294C>T; p.Arg432Ter) associated with non-syndromic hearing loss and a homozygous start-loss variant in *HPDL* (c.3G>C; p.Met1?) associated with NEDSWMA. The coexistence of these variants within the same pedigree resulted in three distinct phenotypic presentations: isolated hearing loss, isolated neurodevelopmental disease, and a blended phenotype in one sibling harboring biallelic pathogenic variants in both genes. These findings illustrate the diagnostic complexity that may arise in highly consanguineous families and emphasize the importance of considering multilocus inheritance when the clinical presentation cannot be explained by a single genetic diagnosis.

The identification of the novel homozygous nonsense variant in *EPS8* expands the mutational spectrum associated with autosomal recessive non-syndromic hearing loss. *EPS8*, encoding Epidermal Growth Factor Receptor Pathway Substrate 8, is one such rare gene implicated in autosomal recessive NSHL. EPS8 is located on chromosome 12p12.3, spans 21 exons, and encodes an 822-amino acid actin-regulatory protein containing pleckstrin homology and SRC homology 3 domains (43). In the inner ear, mechanosensory hair cells detect sound through stereocilia bundles arranged in staircase-like rows of graded height. *EPS8* localizes the tips of the tallest stereocilia, where it contributes to actin polymerization, actin bundling, stereocilia elongation, and hair-cell maturation (44–46). Biallelic pathogenic variants in *EPS8* cause autosomal recessive deafness 102 (DFNB102; MIM #615974), with reported patients typically presenting with congenital or early-onset non-syndromic hearing loss (43,47–49). Previous studies have identified predominantly loss-of-function variants in *EPS8*, supporting biallelic loss of function as the underlying disease mechanism. The novel nonsense variant identified in the present family is predicted to introduce a premature termination codon, likely resulting in nonsense-mediated mRNA decay or production of a truncated protein lacking essential functional domains. Supporting this predicted mechanism, long-read transcriptomic analysis demonstrated the absence of detectable *EPS8* transcripts in both individuals homozygous for the c.1294C>T (p.Arg432Ter) variant, whereas transcript expression was preserved in the heterozygous carrier parents, the unaffected sibling, and the heterozygous sibling affected only by *HPDL*-related disease. Although our transcriptomic analysis did not directly evaluate nonsense-mediated mRNA decay, the absence of detectable *EPS8* transcript is consistent with degradation of mutant transcripts and provides complementary transcript-level evidence supporting a loss-of-function mechanism. The clinical phenotype observed in the affected siblings is consistent with previously reported EPS8-associated hearing loss, further supporting the pathogenicity of this novel variant and expanding the mutational spectrum of *EPS8*. The distribution of previously reported pathogenic variants throughout the coding sequence, together with the segregation pattern observed in our family, further supports the pathogenic role of truncating *EPS8* variants (Figure 3).

Neurodevelopmental disorder with progressive spasticity and brain white matter abnormalities (NEDSWMA; MIM:#619026) is a rare, severe, early-onset neurodegenerative condition caused by biallelic pathogenic variants in the *HPDL* (4-hydroxyphenylpyruvate dioxygenase-like; MIM *618994) gene, located on chromosome 1q34.1 (50). NEDSWA is primarily characterized by developmental delay with motor and cognitive impairment, epilepsy, ophthalmologic abnormalities, and progressive respiratory dysfunction. Recent studies have identified biallelic pathogenic variants in *HPDL* (OMIM *618994) as a cause of neurodevelopmental disorder (51,52). The gene encodes the 4-hydroxyphenylpyruvate dioxygenase-like (HPDL) protein, an enzyme involved in the 4-hydroxymandelate coenzyme Q10 (CoQ10) biosynthetic pathway that is broadly expressed across multiple tissues, with particularly high expression in the central and peripheral nervous systems (52–54). Although the precise biological function of HPDL is not fully understood, accumulating evidence suggests an essential role in mitochondrial metabolism and neuronal homeostasis (55). The twins in our study exhibited a phenotype highly consistent with previously reported cases, including severe global developmental delay, spastic quadriplegia, microcephaly, white matter abnormalities, and profound functional impairment (52,56–58). The homozygous start-loss variant identified in both twins is expected to abolish the canonical translation initiation codon and is therefore predicted to markedly impair protein production. The similarity between our patients and previously reported individuals further strengthens the genotype–phenotype correlation for HPDL-associated disease.

Comprehensive lrWGS was instrumental in resolving the complex molecular basis of disease in this family. Rather than identifying a single genetic diagnosis, genome-wide analysis revealed two independently segregating autosomal recessive disorders responsible for three distinct clinical presentations within the same sibship. This highlights an important advantage of comprehensive genome sequencing over phenotype-driven testing or targeted gene panels, particularly in consanguineous families where multiple rare recessive disorders may coexist. Furthermore, integration of genomic findings with transcriptomic and proteomic analyses provided complementary molecular evidence supporting the pathogenicity of the identified variants and illustrates the value of multi-omics approaches for variant interpretation

To further investigate the molecular consequences of the identified variants, we performed plasma proteomic profiling in all six family members using the SomaScan platform. Distinct protein abundance profiles were observed across family members, with the most pronounced alterations identified in the twins affected by *HPDL*-related neurodevelopmental disease, particularly the individual harboring pathogenic variants in both *EPS8* and *HPDL*. These findings suggest broader perturbation of signaling networks in individuals with dual molecular diagnoses rather than alterations attributable to a single gene. Although these proteomic findings should be interpreted cautiously given the single-family design, they provide complementary molecular evidence supporting the genomic findings and demonstrate the potential utility of integrating proteomic profiling with genomic sequencing in the investigation of complex inherited disorders.

An important consideration when interpreting the transcriptomic and proteomic findings is the tissue-specific biology of EPS8. Although EPS8 is expressed in multiple tissues, its principal biological function is within cochlear sensory hair cells, where it localizes to the tips of the tallest stereocilia and regulates actin dynamics required for stereocilia maturation and mechanotransduction (43,46,59,60). Consequently, transcriptomic and proteomic analyses performed using peripheral blood-derived samples may not fully capture the molecular consequences of the identified variant within the inner ear. Nevertheless, the absence of detectable EPS8 transcripts in the two homozygous individuals provides transcript-level evidence supporting a loss-of-function mechanism, while the plasma proteomic findings offer complementary evidence of altered molecular profiles associated with the underlying genetic defects. These findings should therefore be interpreted as supportive functional evidence rather than direct measures of cochlear pathology.

From a genetic counseling perspective, this family emphasizes the importance of considering multiple independent recurrence risks rather than assuming a single Mendelian disorder. Both parents are heterozygous carriers of pathogenic variants in *EPS8* and *HPDL*, resulting in independent 25% recurrence risks for each autosomal recessive disorder in every pregnancy. Because the two loci segregate independently, there is also a measurable probability of offspring inheriting pathogenic variants in both genes, resulting in a blended phenotype similar to that observed in the female twin. Accurate molecular diagnosis therefore has direct implications for recurrence risk assessment, carrier testing of at-risk relatives, cascade screening within the extended family, and reproductive options including prenatal diagnosis and preimplantation genetic testing for monogenic disorders (PGT-M). In highly consanguineous populations, identification of all segregating pathogenic variants is essential to provide comprehensive counseling and avoid underestimating reproductive risks.

Finally, our findings have broader implications for clinical practice in regions where consanguinity remains common. The presence of multiple recessive disorders within a single pedigree should not be considered exceptional, particularly in populations with extensive endogamy and large extended families. Careful phenotyping, segregation analysis, and comprehensive genomic testing should be considered whenever the clinical presentation is atypical or cannot be fully explained by a single diagnosis. Such an approach improves diagnostic accuracy, facilitates personalized clinical management, and enables informed reproductive decision-making.

This study has several limitations. Functional characterization was limited to transcriptomic and plasma proteomic analyses, and no *in vitro* or *in vivo* experiments were performed to directly investigate the molecular consequences of the identified variants. Furthermore, the transcriptomic and proteomic analyses were performed in a single family, limiting statistical inference and generalizability. Nevertheless, the combination of detailed phenotyping, segregation analysis, long-read whole-genome sequencing, Sanger validation, transcriptomic profiling, and proteomic analysis provides converging evidence supporting the pathogenicity of the identified variants.

In conclusion, we report a consanguineous Pakistani family with dual molecular diagnoses caused by a novel homozygous nonsense variant in *EPS8* and a homozygous start-loss variant in *HPDL*. Our findings expand the mutational spectrum of *EPS8*, further delineate the phenotype associated with *HPDL*-related NEDSWMA, and demonstrate how multilocus pathogenic variation can produce distinct and blended clinical phenotypes within the same family. This study underscores the importance of comprehensive genome sequencing and integrated multi-omics approaches combining long-read whole-genome sequencing, transcriptomics, and proteomics for improving molecular diagnosis, variant interpretation, genetic counseling, and reproductive planning in consanguineous families.

