## Supplementary Figure 1 for "Phenotype Dependent Segregation of a Novel *EPS8* Variant for Hearing Loss and an *HPDL* Variant for Neurodevelopmental Disorders in a Complex Consanguineous Family"

**A. IV-2 (Proband, Homozygous affected)**

| No. | Sample | Gene | Chromosome position | Ref. | Alt. | Result | zygosity |
|---|---|---|---|---|---|---|---|
| 1 | IV-2 | EPS8 | chr12:15650963-15650963 | G | A | A | HOMO |

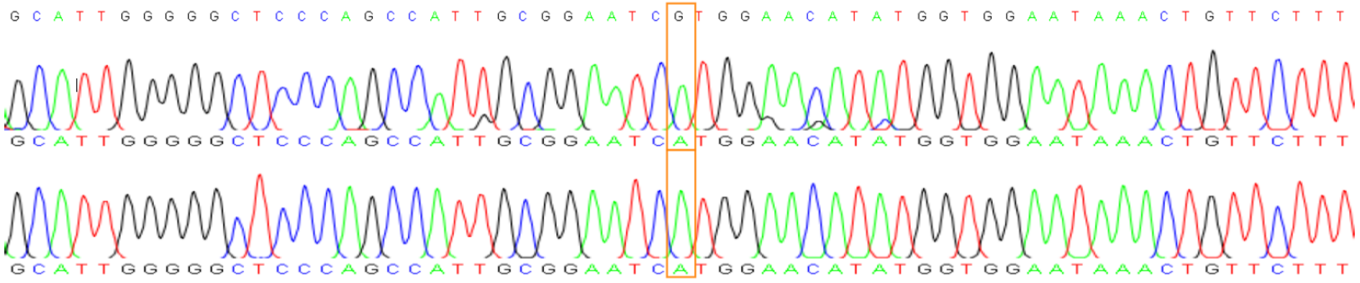


**B. III-1 (Father, Heterozygous carrier)**

| No. | Sample | Gene | Chromosome position | Ref. | Alt. | Result | zygosity |
|---|---|---|---|---|---|---|---|
| 2 | III-1 | EPS8 | chr12:15650963-15650963 | G | A | G/A | HETERO |

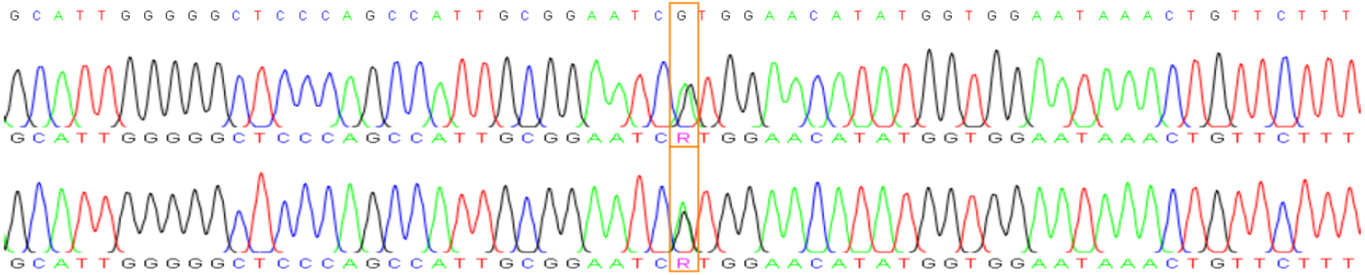


**C. III-2 (Mother, Heterozygous carrier)**

| No. | Sample | Gene | Chromosome position | Ref. | Alt. | Result | zygosity |
|---|---|---|---|---|---|---|---|
| 3 | III-2 | EPS8 | chr12:15650963-15650963 | G | A | G/A | HETERO |

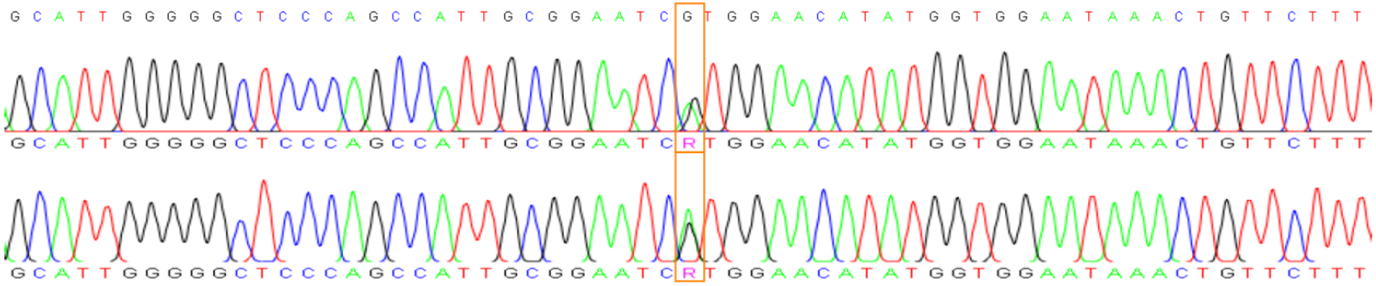
